# Consequences of consumer origin and omnivory on stability in experimental food web modules

**DOI:** 10.1101/589259

**Authors:** Monica Granados, Katie S. Pagnucco, Anthony Ricciardi

**Affiliations:** Department of Biology, McGill University, Montreal, Canada; Redpath Museum and McGill School of Environment, McGill University, Montreal, Canada

**Keywords:** community dynamics, trophic ecology, interaction strength, invasive species, consumer-resource interactions

## Abstract

1. Food web stability, a fundamental characteristic of ecosystems, is influenced by the nature and strength of species interactions. Theory posits that food webs are stabilized by omnivory and disrupted by novel consumers.
2. To test the effects of secondary consumer origin and trophic level on basal resource stability, we constructed crayfish-snail-algae modules using four congeneric species of crayfish (*Faxonius* spp.), two from native populations (*F. propinquus* and *F. virilis*) and two from non-native populations (*F. limosus* and *F. rusticus*). We performed surgical manipulations of crayfish feeding structures to create omnivore food web and predator food chain modules. We compared the temporal stability of these modules using measures of the coefficient of variation of the basal resource (benthic algae).
3. Consistent with theoretical and empirical predictions, food web modules with omnivory had the lowest coefficient of variation. However, contrary to prediction, we did not find consistently higher coefficients of variation in modules with non-native species. Rather, across species, we found the lowest coefficient of variation in modules with one of the non-native species (*F. rusticus*) and one native species (*F. virilis*), owing to stronger interactions between these crayfish species and their snail and algal food resources.
4. The results suggest that omnivory is indeed stabilizing and that very weak interactions or very low attack rates of the consumer on the basal resource can be unstable. Thus, we demonstrate that omnivores may have different impacts than predators when introduced into a novel ecosystem, differences that can supersede the effect of species identity.

## Introduction

Food web stability, here defined as temporal constancy, is a fundamental characteristic of ecosystems (Worm & Duffy, 2003) that can be profoundly affected by the presence of omnivores – organisms that feed on more than one trophic level (Pimm & Lawton, 1978; Pimm, 1982). Omnivory is common in food webs across a broad range of habitats, including freshwater systems (Thompson et al., 2007; Thompson, Dunne, & Woodward, 2012; Wootton, 2017). Omnivores reduce the strength of consumer-resource links by shunting some of the energy up the omnivore-resource pathway and away from the consumer-resource pathway (McCann, Hastings, & Huxel, 1998). A form of omnivory that has been the focus of theoretical and empirical investigations on stability is intraguild predation, where an omnivore feeds on an intermediate consumer in addition to one of the prey’s resources (Polis, Myers, & Holt, 1989; Holt & Polis, 1997). Through predation (Fig. 1A), the omnivore increases the mortality of the common resource, thereby preventing the latter from experiencing overshoot population dynamics (McCann, 2012); for example, if the population of the intermediate consumer suddenly declines, omnivore predation can prevent the population of the common resource from increasing in response.

**Figure 1.**
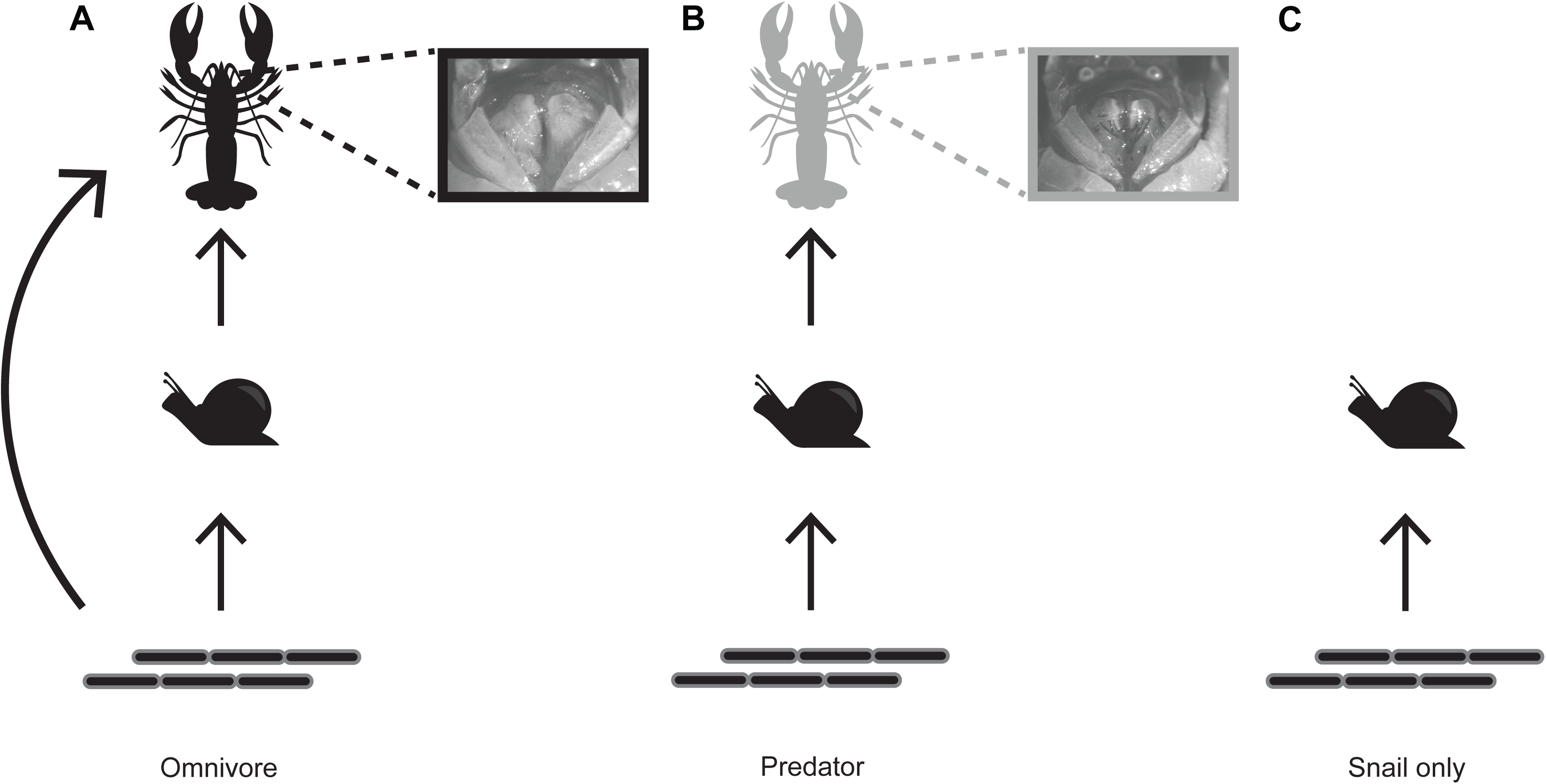
Modules used in the experiment. **A.** Omnivore food web module. Omnivore crayfish (black) both consumed and competed with snails for a common resource (benthic algae). **B.** Predator food chain module. Predator-converted crayfish (grey) consumed only snails, which in turn consumed benthic algae. Insets depict the removal of setae in the predator treatment to prevent the consumption of benthic algae. **C.** Snail-only, consumer-resource interaction, treatment.

Experimental studies of the dynamics of simple three- or four-species food webs with and without omnivores have revealed that omnivory is stabilizing (Lawler & Morin, 1993; Morin & Lawler, 1995). One of the few direct experimental tests of the effect of omnivory on food web stability was conducted on arthropod assemblages by Fagan (1997), who found that a high degree of omnivory stabilized community dynamics following disturbance; however, the omnivore and predator species used in the experiment comprised different genera and, consequently, the effects of omnivory on community stability were confounded by potential species effects.

There are also reasons to expect consumer origin to influence food web stability. Non-native consumers generally have stronger negative effects than trophically-similar natives on native prey populations (Salo et al., 2007; Paolucci, MacIsaac, & Ricciardi, 2013). These effects are thought to result, at least in part, from prey naïveté wherein prey have not had selective pressures to adapt defences to novel predator traits (Cox & Lima, 2006). Moreover, non-native populations of predators and consumers tend to have higher resource consumption rates and can thus exert greater impacts on food resources (Bollache et al., 2008; Morrison & Hay, 2011; Dick et al., 2013; Iacarella et al., 2015). Such mechanisms can produce strong interactions that are destabilizing (McCann, Hastings & Huxel, 1998). Non-native species could also potentially disrupt food webs by being stronger interactors than trophically similar natives, or by eliminating other species, consequently increasing the average interaction strength within a food web (Barrios-O’Neill et al., 2014).

Crayfishes are among the most common omnivores in freshwater ecosystems and their activities can structure littoral food webs (Olsen et al., 1991; Dorn & Wodjak, 2004; Nilsson et al., 2012; Twardochleb, Olden & Larson, 2013). Non-native crayfishes have been widely introduced into lakes and rivers, where they can replace native crayfishes (Lodge, 1987; Lodge et al., 2000), significantly reduce macroinvertebrate grazer densities (Dorn, 2013) so as to cause trophic cascades (Charlebois & Lamberti, 1996), and trigger other complex indirect effects that lead to changes in the structure of communities and food webs (Taylor & Redmer, 1996, Nystrom et al., 1996; Lodge et al., 2000; Wilson et al., 2004; Twardochleb et al., 2013). Recognizing crayfishes as potentially valuable model organisms for studying food web dynamics, we used individuals from two native and two non-native populations, and surgically manipulated their mouthparts to alter their trophic guild, with the aim of testing the effects of secondary consumer origin and trophic level on stability in an experimental tri-trophic food web. We hypothesized that 1) omnivory in the food web will mute oscillations in the basal resource and result in greater stability – indicated by a lower coefficient of variation in the resource; and 2) strong interactions involving non-native species will produce a higher coefficient of variation in the resource compared to native crayfishes, owing to extinction of the primary consumer and concomitant release of top-down control on the resource.

## Materials and methods

### Experimental design

We created modules of a crayfish-snail-algae food web using four congeneric species of crayfish (*Faxonius* spp.) and a snail-only consumer-resource interaction for comparison (Fig. 1). Two crayfish species were collected from populations in their native range (the northern clearwater crayfish *Faxonius propinquus* and the virile crayfish *F. virilis*) and two other species were from non-native populations (the spinycheek crayfish *F. limosus*, and the rusty crayfish *F. rusticus*) (see Supplementary Information for locations). Each of these species includes snails in its natural diet (Crowl & Covich, 1990; Rosenthal, Stevens, & Lodge, 2006; Twardochleb et al., 2013; A.R., pers. obs.).

The experiment was conducted in 36 outdoor freshwater mesocosms (114-L plastic containers, 81 × 51.4 × 44.5 cm) located at McGill University (Montreal, Quebec). Each mesocosm contained 4 L of gravel sediment to foster natural biogeochemical cycling processes and, on 15 July 2013, they were filled with 64 L of dechlorinated tap water. Benthic algae were allowed to grow on tiles placed at the bottom of mesocosms for 21 days prior to the start of experiments. Subsequently added to each mesocosm were 70 snails (*Physella* sp.), which acted as primary consumers of benthic algae. Finally, each mesocosm received a single crayfish from one of the four species (*F. propinquus, F. virilis, F. limosus* and *F. rusticus*), which behaved as either an omnivore (consumed benthic algae) or a predator (did not consume benthic algae), depending on surgical manipulation (procedures described below). Each food web module × species combination and the snail-only module were repeated four times, for a total of 36 mesocosms.

All mesocosms were arranged adjacent to each other in a single row, across which modules were distributed randomly. A refuge (PVC pipe, 10 cm length × 5 cm diameter) was also added to each mesocosm to reduce crayfish stress. Eight 10 cm × 10 cm tiles that were divided into quadrats were attached to the bottom of each mesocosm using magnets to keep them stationary during the experiment. The tiles were used as substrate on which the algae would grow, and from which we would collect algal samples for analysis. Mesocosms were covered with 2 mm^2^ vinyl mesh to reduce colonization by macroinvertebrates, to minimize diurnal temperature variations, and to prevent crayfish from escaping.

### Procedures for predator conversion

Although crayfish remove benthic algae less efficiently than snails do (Luttenton, Horgan, & Lodge, 1998), they feed on both vegetation and animals to a sufficient degree to be classified as omnivores and they exhibit a specificity in feeding structures for different resources (Holdich, 2002). We created phylogenetically-equivalent ‘predators’ to compare against omnivores by manipulating crayfish surgically to prevent them from consuming algae and thus rendering them a default predator. The transformation of crayfish from omnivores to predators was achieved by altering their “filter proper”, which is comprised of the acuminate setae on the 1^st^ maxilliped and maxillae (Budd, Lewis, & Tracey, 1978; Holdich, 2002). Setae from the 1^st^ to 3^rd^ maxillipeds, maxilla, maxillule and mandible were removed under a microscope using microdissection scissors while crayfish were under anesthesia (clove oil at 1 ml/L). Crayfish selected as omnivores were also anesthetised and placed under a microscope for the same duration as the full predator conversion procedure; this was intended to reduce any manipulation effects on subsequent crayfish behaviour. Dissections were performed from 31 July to 3 August 2013 (the date of the procedure was randomised across species), after which the crayfish were kept in separate tanks during the recovery period prior to the beginning of the experiments. The manipulation of arthropod mouthparts has been used to control predation in experiments (e.g., Schmitz et al., 1997; Nelson, Matthews, & Rosenheim, 2004); however, in these previous studies mouthparts were altered to prevent consumption of all prey, whereas in the present study mouthparts were manipulated to restrict consumption to certain resources.

### Sampling benthic algal density and snail abundance

The experimental period lasted 61 days (5 August – 6 October 2013). Benthic algal density was sampled every second day over the experimental period. On each sampling day, algae were scraped from a single quadrat within a single tile from the bottom of each mesocosm; the quadrat was chosen randomly for each mesocosm and sampled once over the experimental period. The sample was then added to 30-mL of dechlorinated tap water, and the concentration of chlorophyll-*a* in each sample was determined using fluorometry (FluoroProbe, bbe-Moldaenke, Kiel, Germany). Thus, chlorophyll-*a* concentration was used as a proxy measure of benthic algal density. Data from eight tiles were collected from a total of four replicates for all crayfish species for each module, except for the *F. rusticus* predator food chain module, where data was collected from only three mesocosms owing to crayfish mortality.

### Statistical analyses

To measure snail mortality (including loss due to crayfish predation) across treatments, the day at which 75% of snails were lost in each mesocosm (LD75) was estimated by fitting a binomial model with a logit link function to the snail density time series data. Here we use LD75 in an analogous way to toxicity studies: as the LD75 decreases, the rate of mortality increases. Treatments with a lower LD75 suffered higher mortality (a more rapid onset of 75% mortality) than higher LD75 values. A two-way analysis of variance (ANOVA) was then performed on mean LD75 using food web module and species as fixed factors.

Owing to the high degree of spatial variation within the mesocosms, for each tile in a mesocosm, we calculated the net algal density change, i.e.

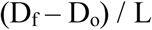

where D_f_ = density on the last day that tile was sampled, D_o_ = density on the first day that tile was sampled, and L = length of sampling in days. We then averaged across the 8 values per mesocosm to obtain the mean net algal density change for each mesocosm. A two-way analysis of variance (ANOVA) was performed on the mesocosm net algal density change using food web module and species as fixed factors.

To assess the stability of benthic algal density, analyses were focused on temporal stability using measures of the coefficient of variation within tiles across time (Pimm, 1991; Tilman et al., 2006; Ives & Carpenter, 2007). Coefficient of variation, equal to the standard deviation divided by the mean, is a scale-independent measure of variability that is used in ecological studies (Haddad et al., 2011; Schindler et al., 2010; Howeth & Leibold, 2010; Kratina et al., 2012). A two-way analysis of variance (ANOVA) was then performed on the mean coefficient of variation for each mesocosm using food web module and species as fixed factors. All statistics and figures were performed using R (R Core Team, 2017).

## Results

Supporting our first hypothesis, algal densities in the omnivore food web module were less variable than in the predator food chain module (Fig. 2, ANOVA, Tukey HSD, *P*=0.0429). However, neither the omnivore nor the predator modules differed from the snail-only module in terms of variability (ANOVA, Tukey HSD, *P*=0.5441 and *P*=0.8345, respectively). The coefficient of variation differed between *F. rusticus* and *F. limosus* (Fig. 2, ANOVA, Tukey HSD, *P* = 0.0023) and between *F. virilis* and *F. limosus* (Tukey HSD, *P* = 0.0213), but not between other species pairs (Table S1 in Supporting Information).

**Figure 2.**
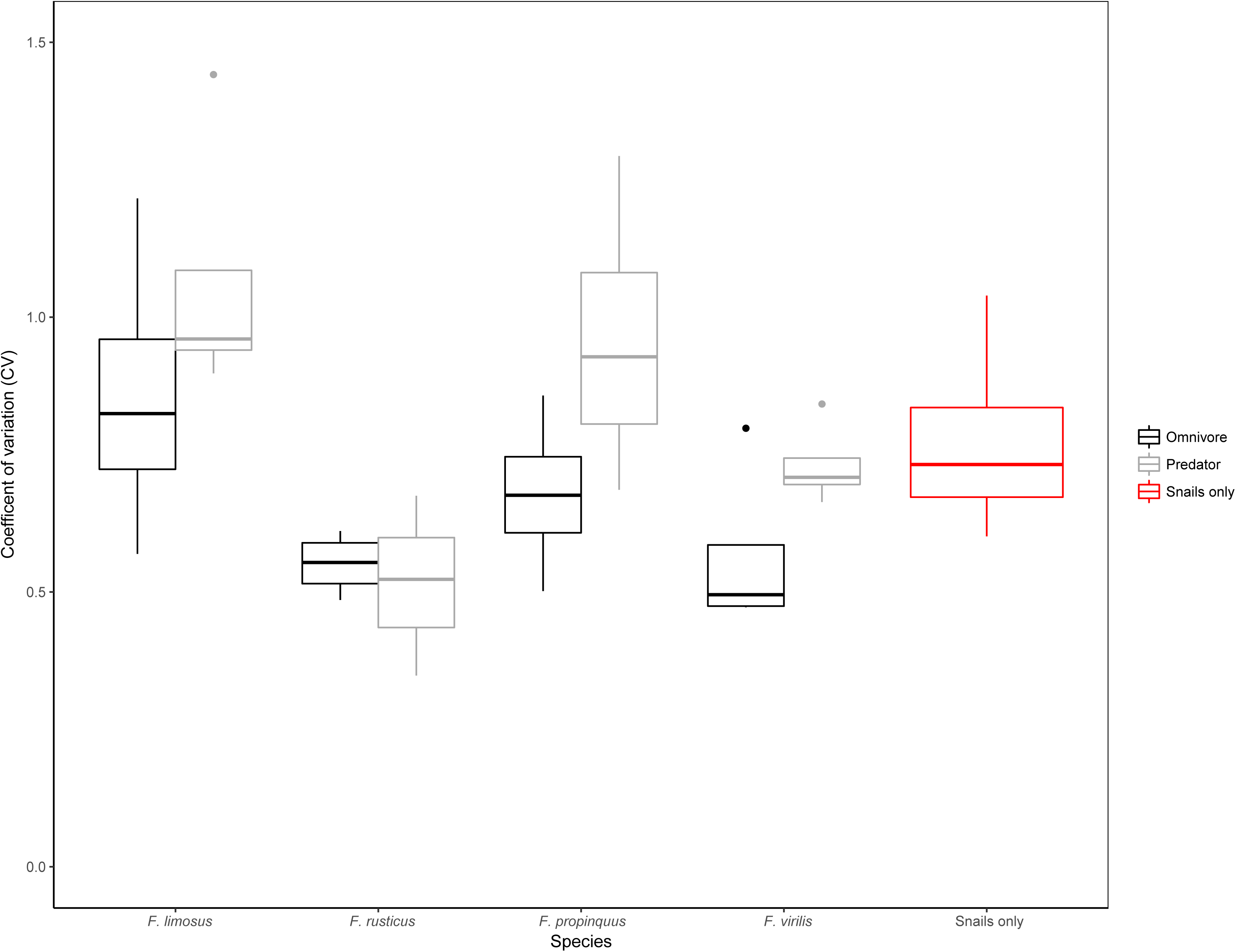
Median coefficient of variation (standard deviation/mean) for benthic algal density across the experiment for each module. The lower and upper hinges correspond to the first and third quartiles (the 25th and 75th percentiles). The coefficient of variation in the predator food chain module is significantly higher than the omnivore food web (ANOVA, Tukey HSD, *P* = 0.0429) but not the snail-only modules (ANOVA, Tukey HSD, *P* = 0.8345). Crayfish are grouped by origin on the x-axis. *F. limosus* and *F. rusticus* are non-native, while *F. propinquus* and *F. virilis* are native species. Variation tended to be higher in association with omnivores than predators for each of the crayfish species except, *F. rusticus*, which regenerated its filter proper such that the predator treatment was ineffective (see Discussion).

The rate of snail mortality was greater in the omnivore food web module than the predator food chain and snail-only modules (ANOVA, Tukey HSD, *P* = 0.0247, *P*<0.001, Fig. 3A; see Fig. S1 in Supporting Information, for estimated snail abundances over time in each module). It was also greater in the predator food web module than in the snail-only module (ANOVA, Tukey HSD, *P* = 0.001). Contrary to our second hypothesis, consumer origin had no consistent effect on snail mortality and variability in algal densities. Snails suffered a higher rate of mortality in the presence of non-native *F. rusticus* compared to either non-native *F. limosus* or native *F. propinquus* (ANOVA, Tukey HSD, *P* =0.0010, *P=*0.0331, Fig. 3A, Table S2 in Supporting Information), and in the presence of native *F. virilis* compared to non-native *F. limosus* (ANOVA, Tukey Test, *P* =0.0083). Evidence of predation was provided by fragments of crushed shells found only in the presence of crayfish. Snail mortality in the snail-only treatment was presumed to result from intraspecific competition.

**Figure 3.**
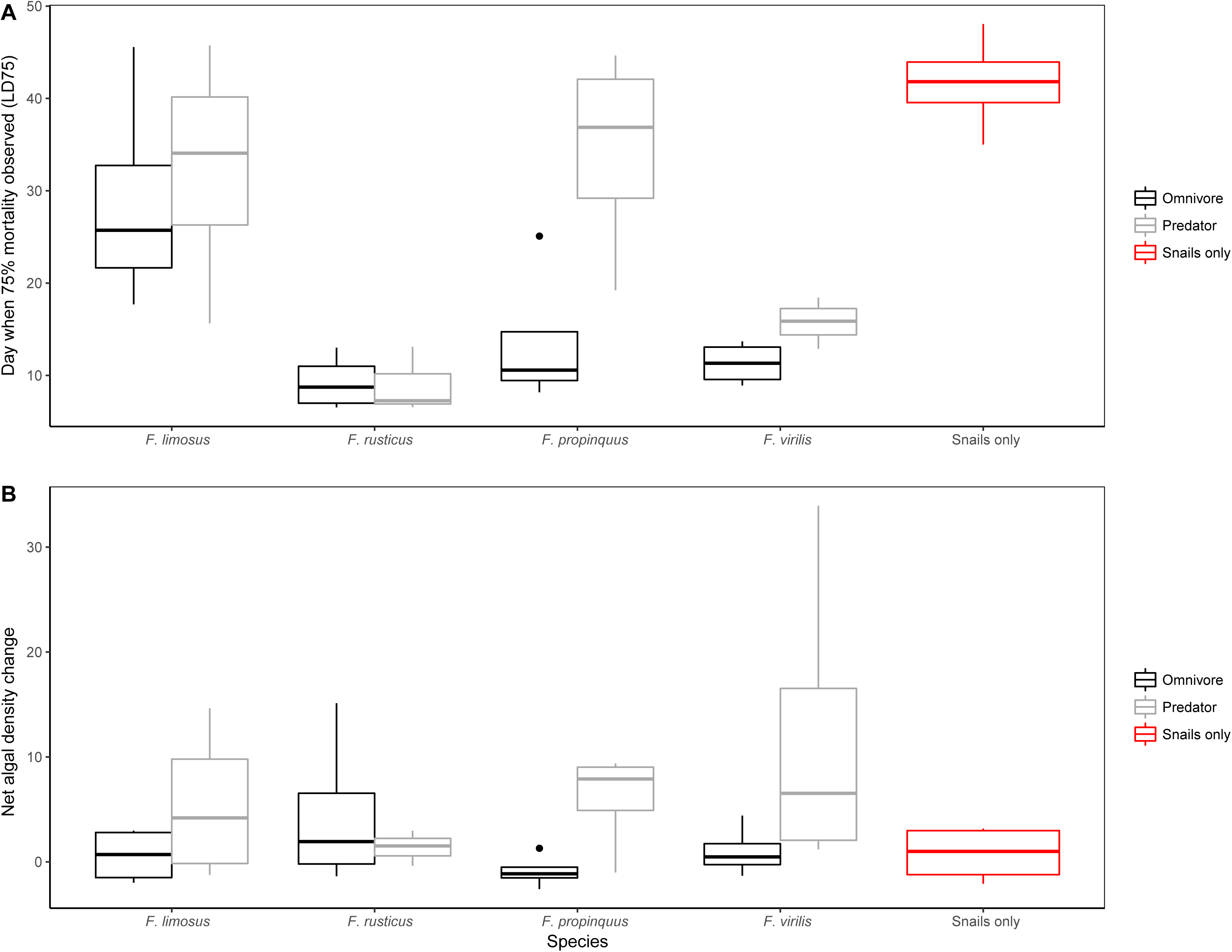
**A.** Median day each species consumed 75% of the snails available in each mesocosm (LD75) for each module. The lower and upper hinges correspond to the first and third quartiles (the 25th and 75th percentiles). Across all species, omnivores consumed 75% of snails in the mesocosm (i.e., reached LD75) faster than predators (ANOVA, Tukey HSD, *P* = 0.0416) but not the snail-only module (ANOVA, Tukey HSD, *P* = 0. 7067). **B.** Mean net algal density change for each food web module and species. The lower and upper hinges correspond to the first and third quartiles (the 25th and 75th percentiles). Crayfish are grouped by origin on the x-axis. *F. limosus* and *F. rusticus* are non-native, while *F. propinquus* and *F. virilis* are native species.

Changes in algal densities tended to be greater in the predator food web module compared with the omnivore food web module (Fig. 3B, ANOVA, Tukey HSD, P = 0.0957), but not the snail-only module (ANOVA, Tukey HSD, P = 0.9897). There was no difference in net algal density change between the predator and the snail-only modules (ANOVA, Tukey HSD, P = 0.2921). Nor was there a significant difference in net algal density change among species (Table S3 in Supporting Information).

## Discussion

Theory suggests that omnivory increases stability by weakening coupling strengths that otherwise create large oscillations in organismal populations (McCann & Hastings, 1997; McCann et al., 1998; McCauley, Jenkins & Quintana-Ascencio, 2013). Two previous studies found empirical evidence of omnivory increasing stability (Fagan, 1997; Holyoak & Sachdev, 1998), but ours provides the first phylogenetically-controlled test of this phenomenon. Consistent with our first hypothesis, we found that the coefficient of variation was lower in food web modules with omnivores. Here, the interaction between the omnivore and benthic algae reduced the energy flux between the snail and benthic algae. Although the predation rate on snails was greater in the omnivore food web module than in the predator food web module, all treatments reached the LD75 – and thus benthic algae were released from predation – within the time frame of our experiment. We posit that the predator food web modules reduced snail abundances at a slower rate owing to latent effects of the removal of the crayfish’s filter proper. Nevertheless, the depletion of snails in all treatments created the potential for benthic algal densities to increase rapidly. However, because the omnivorous crayfish could consume benthic algae, it prevented the algae from growing unchecked after snails were reduced. In the predator food chain module, the removal of snails resulted in a significant increase in the net algal density change and a higher coefficient of variation. These results are consistent with theoretical predictions that if a consumer-resource interaction is excitable or shows oscillations of any sort, removing biomass can stabilize the interaction (McCann, 2012). Here, the snail-benthic algae interaction shows oscillatory potential: as the snail population decreases, the benthic algae population increases. In the omnivory modules, however, benthic algae were removed, thereby weakening the relative coupling strength of the snail-benthic algae interaction. Thus, our study demonstrated the ability of an omnivore to increase temporal stability in the resource compared to a predator that shares the entire suite of species traits except for the adjustment of mouthparts.

Our results did not support our prediction of the effect of species origin on stability. High resource consumption rates and efficient prey handling times are linked to the invasion success and impact potential of crayfishes (Haddaway et al., 2012; Taylor & Dunn, 2018). Therefore, we expected that crayfishes from non-native populations would reduce stability through higher snail consumption rates compared with those from native populations (Barrios-O’Neill et al., 2014). Snail mortality was higher in the presence of non-native *F. rusticus* than with native *F. propinquus* and non-native *F. limosus*, but not in comparison with native *F. virilis.* Owing to the lack of consistently higher snail consumption in non-native species, stability was not affected by crayfish origin in our study. We propose three explanations for this. First, each of the four crayfish species used in our experiment has a history of invasion and ecological impacts beyond its native range (Wilson et al., 2004; Rosenthal et al., 2006; Twardochleb et al., 2013). Both *F. limosus* and *F. rusticus* have extensive invasion histories and have caused significant impacts on recipient communities in North America and Europe (Olsen et al., 1991; Kozák et al., 2007; Hirsch, 2009; Nilsson et al., 2012). The virile crayfish *F. virilis* occurs naturally over a large area of the USA and Canada (encompassing the Great Lakes-St Lawrence River basin and extending to the continental divide) and has been introduced to other regions of North America (Hobbs, Jass & Huner, 1989; Phillips, Vinebrooke & Turner, 2009; Larson et al., 2010) and Europe (Ahern, England, & Ellis, 2008). The northern crayfish, *F. propinquus*, has a relatively limited invasion history, but has also caused impacts in recipient systems (Hill & Lodge, 1999; Rosenthal, Stevens, & Lodge, 2006). Traits contributing to trophic impacts might be conserved across conspecific populations, such that biogeographic origin is not as influential as other environmental factors in this context. Indeed, it has been suggested that non-native crayfish identity is less important than extrinsic characteristics of invaded ecosystems in determining their impact (Twardochleb et al., 2013), but testing this hypothesis requires multi-species and multi-site comparisons. Second, traits found to be most important for invasion success in crayfish (i.e., aggression, boldness, fecundity; Lindqvist & Huner, 1999; Gherardi & Cioni, 2004; Hudina & Hock, 2012) might not be relevant to our experiment, although they are correlated with prey consumption rates (Pintor et al., 2008). Third, the prey species (*Physella* sp.) used in our experimental food webs has evolutionary experience with crayfishes, including *Faxonius* spp., such that it can respond to their cues (Klose et al., 2011) and, thus, it is not as naïve to the consumers used in our experiment as it would be to a novel predator/omnivore archetype (Cox & Lima, 2006).

We did not directly test the efficacy of our surgical manipulation. However, the contrasts between the omnivore and predator effects on algal density (Fig. 2B) suggest a differential efficiency in removing algae. It is interesting that, in this regard, the least difference was exhibited by *F. rusticus*, which is the most successful invader among the species used here (Wilson et al., 2004). In this species we observed the regeneration of the filter proper within 60 days, whereas no evidence of regeneration was observed in the other species within the experimental time frame. The regeneration of its filter proper might reflect a capacity for rapid growth and plasticity, which are putative traits of highly successful invaders (Crispo, 2010; Sargent & Lodge, 2014).

Although differential snail consumption across crayfish species did not produce differences in benthic algal densities, there were differences in the coefficient of variation. Stronger interaction strengths involving *F. rusticus* and *F. virilis* produced lower coefficients of variation in the basal resource than did *F. limosus* – the species with the longest LD75 or, ostensibly, the weakest interaction strength between crayfish, snails and benthic algae. Our results suggest that although weak to intermediate interaction strengths are stabilizing in food webs, very weak interactions are destabilizing. This is consistent with theoretical results in Lotka-Volterra models, which indicate that when the attack rates are very weak the dominant eigenvalue is positive or unstable (McCann, 2012).

Taken together, these results are consistent with previous empirical work showing omnivory is stabilizing, and that the trophic position of the species – but not its origin – has an important effect on the stability of the resource population. Our study suggests that, when omnivory is weak to intermediate, non-native omnivores can also potentially stabilize the consumer-resource interactions in comparison to predators. This merits further examination, given that freshwater food webs are subject to an increasing number of introduced species, many of which are omnivores (Wootton, 2017).

## Data Accessibility

The complete R script for the analyses performed in the paper and associated data can be found online archived on Zenodo at the following URL: https://zenodo.org/record/1341885

## Acknowledgements

We thank Rowshyra Castañeda, Sian Kou-Giesbrecht, and François Vincent for their assistance with the experiment, Brie Edwards, Keith Somers and Ron Ingram for their support in the collection of crayfish. This study was funded by the National Sciences and Engineering Research Council of Canada and the Canadian Aquatic Invasive Species Network.

## Supporting Information

The following Supporting Information is available for this article online.

### Crayfish and snail collection

**Figure S1.** Mean snail abundances, averaged across species x trophic level replicates, across time for species in the experiment (A) and the snail-only module (B). The smoothing line is a locally weighted scatterplot smoothing (LOESS) regression. Crayfish are grouped by origin on the x-axis. *F. limosus* and *F. rusticus* are non-native, while *F. propinquus* and *F. virilis* are native species.

**Table S1.** Tukey HSD results from a two-way ANOVA with food web module and species as fixed factors for the algal coefficient of variation response variable.

**Table S1.** Tukey HSD results from a two-way ANOVA with food web module and species as fixed factors for the LD75 response variable.

**Table S2.** Tukey HSD results from a two-way ANOVA with food web module and species as fixed factors for the net algal density change response variable.

