## Supporting Information for "Consequences of consumer origin and omnivory on stability in experimental food web modules"

### Crayfish collection

Eight *F. rusticus* (mean carapace length ± 1SE, 25.26 ± 0.43 mm) were collected from Little Rouge River in Ontario (43°50’8.8794”N, 79°11’37.5354”W) on 29 July 2013, eight *F. propinquus* (26.68 ± 0.58 mm) and eight *F. virilis* (30.48 ± 0.85 mm) were collected from Blue Chalk Lake in Ontario (45°12’1.764”N, 78°56’50.352”W) on 30 July 2013, and eight *F. limosus* (21.66 ± 0.69 mm) were collected from the St. Lawrence River near Parc René-Lévesque at Lachine, Quebec (45°25’40.5624”N, 73°40’41.1882”W) on 2 August 2013. Snails (*Physella* spp.) were also collected from the St. Lawrence River near Parc René-Lévesque from 31 July – 3 August 2013.

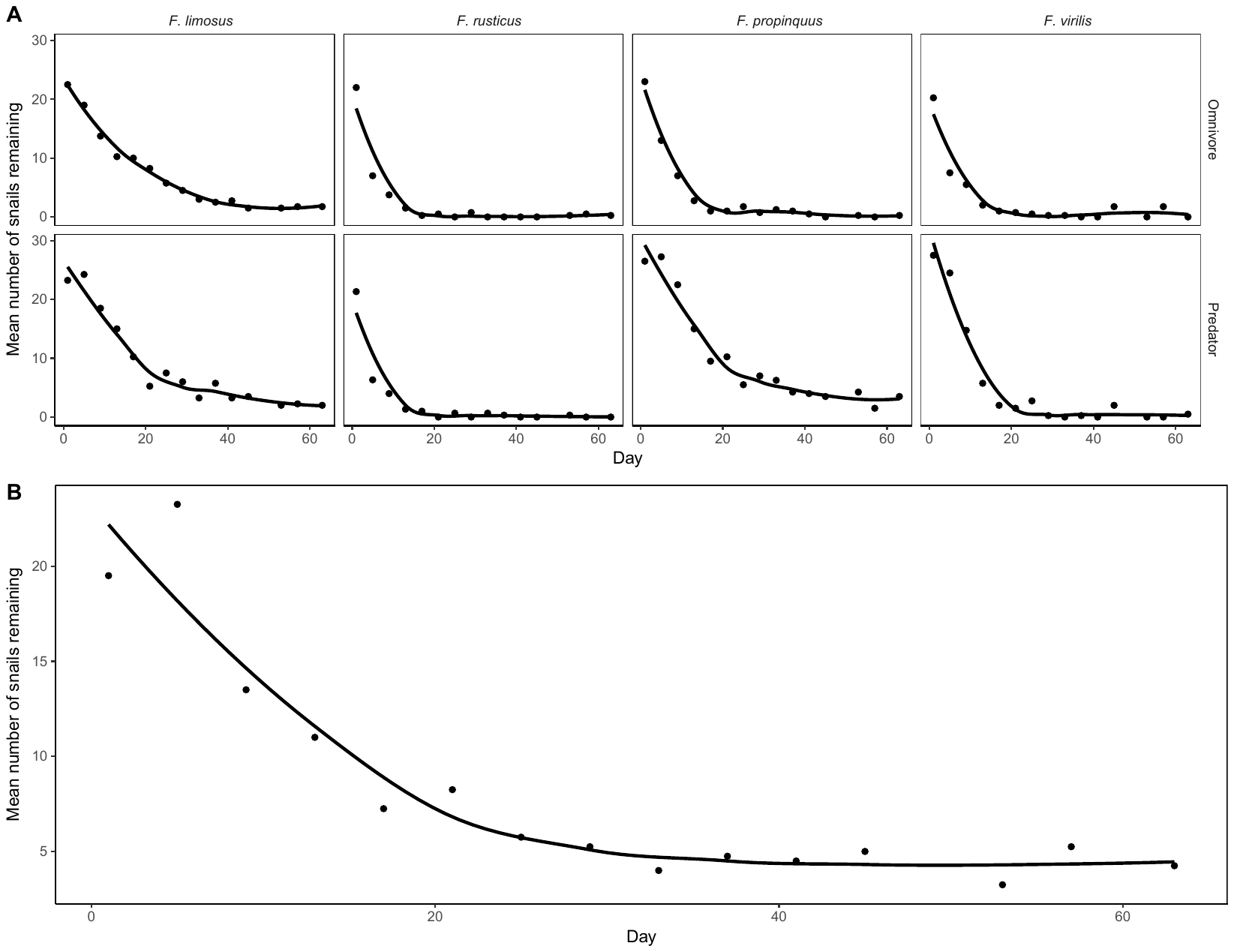

Figure S1. Mean snail abundances, averaged across species x trophic level replicates, across time for each A. species in the experiment. B. the snail-only food web module. The smoothing line is a locally weighted scatterplot smoothing (LOESS) regression.

| Table S1. Tukey HSD results from a two-way ANOVA with food web module and species as fixed factors for the algal coefficient of variation response variable. | | | | | |
| --- | --- | --- | --- | --- | --- |
| Species | *F. limosus* | *F. rusticus* | *F. propinquus* | *F. virilis* | Snail-only |
| *F. limosus* | - | 0.0023 | 0.5649 | 0.0213 | 0.3854 |
| *F. rusticus* |  | - | 0.0737 | 0.8464 | 0.4524 |
| *F. propinquus* |  |  | - | 0.3984 | 0.9763 |
| *F. virilis* |  |  |  | - | 0.9014 |
| Snail-only |  |  |  |  | - |
| Table S2. Tukey HSD results from a two-way ANOVA with food web module and species as fixed factors for the LD75 response variable. | | | | | |
| Species | *F. limosus* | *F. rusticus* | *F. propinquus* | *F. virilis* | Snail-only |
| *F. limosus* | - | 0.0010 | 0.6328 | 0.0083 | 0.3135 |
| *F. rusticus* |  | - | 0.0331 | 0.8899 | <0.001 |
| *F. propinquus* |  |  | - | 0.1915 | 0.0320 |
| *F. virilis* |  |  |  | - | <0.001 |
| Snail-only |  |  |  |  | - |
| Table S3. Tukey HSD results from a two-way ANOVA with food web module and species as fixed factors for the net algal density change response variable. | | | | | |
| Species | *F. limosus* | *F. rusticus* | *F. propinquus* | *F. virilis* | Snail-only |
| *F. limosus* | - | 0.9999 | 0.9999 | 0.8311 | 0.9995 |
| *F. rusticus* |  | - | 0.9989 | 0.9020 | 1.0000 |
| *F. propinquus* |  |  | - | 0.7617 | 0.9974 |
| *F. virilis* |  |  |  | - | 0.9675 |
| Snail-only |  |  |  |  | - |
